# Individualized functional connectivity markers associated with motor and mood symptoms of Parkinson’s disease

**DOI:** 10.1101/2024.01.31.578238

**Authors:** Louisa Dahmani, Yan Bai, Wei Zhang, Jianxun Ren, Shiyi Li, Qingyu Hu, Xiaoxuan Fu, Jianjun Ma, Wei Wei, Meiyun Wang, Hesheng Liu, Danhong Wang

## Abstract

Parkinson’s disease (PD) is a complex neurological disorder characterized by many motor and non-motor symptoms. While most studies focus on the motor symptoms of the disease, it is important to identify markers that underlie different facets of the disease. In this case-control study, we sought to discover reliable, individualized functional connectivity markers associated with both motor and mood symptoms of PD. Using functional MRI, we extensively sampled 166 patients with PD (64 women, 102 men; mean age=61.8 years, SD=7.81) and 51 healthy control participants (32 women, 19 men; mean age=55.68 years, SD=7.62). We found that a model consisting of 44 functional connections predicted both motor (UPDRS-III: Pearson *r*=0.21, FDR-adjusted *p*=0.006) and mood symptoms (HAMD: Pearson *r*=0.23, FDR-adjusted *p*=0.006; HAMA: Pearson *r*=0.21, FDR-adjusted *p*=0.006). Two sets of connections contributed differentially to these predictions. Between-network connections, mainly connecting the sensorimotor and visual large-scale functional networks, substantially contributed to the prediction of motor measures, while within-network connections in the insula and sensorimotor network contributed more so to mood prediction. The middle to posterior insula region played a particularly important role in predicting depression and anxiety scores. We successfully replicated and generalized our findings in two independent PD datasets. Taken together, our findings indicate that sensorimotor and visual network markers are indicative of PD brain pathology, and that distinct subsets of markers are associated with motor and mood symptoms of PD.

## INTRODUCTION

Parkinson’s disease (PD) is a neurologic disorder characterized by multiple different symptoms throughout its course. In the prodromal phase, the disease is marked by non-motor symptoms, such as depression, olfactory dysfunction, sleep disturbances, constipation, and cognitive impairment. Then, the classical motor symptoms associated with PD appear, and are accompanied by worsening non-motor features. To provide a comprehensive picture of the pathophysiology of PD, it is crucial to identify markers that underlie the various features of the disease—both motor and non-motor.

Resting-state functional connectivity (rs-FC), as measured with functional MRI (fMRI), offers a promising avenue for this endeavor. The diversity of PD symptoms indicates that several brain regions and or/networks are disrupted by the disease, and, indeed, there is evidence that rs-FC is directly disrupted by PD neuropathology^1–3^. Studies that investigated large-scale brain networks in PD reported widespread abnormal functional connectivity across networks^1, 4–13^. In some cases, network disruptions were associated with motor and/or non-motor symptoms^4–9, 13, 14^. However, while various network dysfunctions have been identified, replicability has seldom been pursued^15^.

This endeavor is marred with challenges, which include heterogeneity in symptom presentation, a lack of individualized brain-based measurements, and low quality and quantity of imaging data, factors ultimately leading to low replicability and generalizability of identified markers. In particular, a lack of individualized brain measurements entails that any identified brain marker is an approximation based on an atlas or a group average, which tends to translate poorly to other cohorts and to individual patients. We have demonstrated that the functional organization of the brain varies widely from individual to individual, especially when it comes to the association cortices^16, 17^, and that the use of individual-based methods markedly improves the detection of brain-behavior relationships as compared to group-average methods^18–25^. Additionally, clinical MRI studies generally acquire small quantities of imaging data per patient, while achieving reliable functional measures at the individual level requires significant amounts of data.

Here, we sought to identify reliable markers of motor and non-motor symptoms of PD. We addressed the aforementioned challenges by acquiring extensive fMRI data (31min 10sec per subject) in a large dataset of patients with PD and applying individual-based methods to obtain precise, idiosyncratic fMRI measurements of functional connectivity. Importantly, we investigated the replicability and generalizability of our findings by using independent datasets.

## RESULTS

### Participants

Four patients with PD from the main cohort did not complete the study and had incomplete or no MRI scanning, and 10 patients were excluded due to excessive head motion. Therefore, 166 patients were included in the study (64 women, 102 men; mean age=61.8 years, SD=7.81; see demographic and clinical information in Table 1). A total of 51 healthy control participants were included in the study (32 women, 19 men; mean age=55.68 years, SD=7.62; Table 1). The control group was significantly different from the PD group in terms of age (independent samples t-test, *t*(215)=4.92, *p*<0.01), sex (Chi square test Χ^2^=8.30, *p*<0.01), and education (independent samples t-test, *t*(215)=-2.37, *p*=0.01, 2-tailed). The average head motion was slightly lower in the PD group (mean=0.10, SD=0.04) than in the control group (mean=0.12, SD=0.03) (independent samples t-test, t=-2.92, p=0.004).

**Table 1.** Demographic and clinical information.

|  | PD | HC |
| --- | --- | --- |
| Demographic information |  |  |
| Sample size | 166 | 51 |
| Sex | 64 F / 102 M | 32 F / 19 M |
| Age | 61.80±7.81 | 55.68±7.62 |
| Years of education | 8.60±3.74 | 10.01±3.71 |
| Clinical information |  |  |
| Disease duration in years | 5.82±4.06 | - |
| UPDRS-III | 38.36±19.85 | - |
| Hoehn-Yahr | 2.35±0.90 | - |
| Tremor | 8.45±5.23 | - |
| Bradykinesia | 9.93±6.24 | - |
| Rigidity | 17.64±10.73 | - |
| Gait | 2.30±1.77 | - |
| Upper extremities | 16.37±8.77 | - |
| Lower extremities | 9.37±7.01 | - |
| HAMD | 10.78±5.78 | - |
| HAMA | 10.72±6.22 | - |
| MMSE | 25.57±4.15 | - |
| Means ± standard deviations are reported.<br>HC=Healthy control; PD=Parkinson's disease |  |  |

### Reliable connectivity markers of PD

We sought to identify the functional connections exhibiting consistent, significant differences in FC strength between the PD and control groups (see Methods and depicted steps in Fig. 1A). Several functional connections were consistently different between the PD discovery subset (n=112) and control groups. A total of 7,001 connections, whose FC strengths were significantly different from 0 in the control group (one-sample t-tests, *p*’s<0.05), were used as candidate features; 1,181 were significantly different between the two groups (independent samples t-tests, FDR *q*=0.05), of which 44 were consistent (Fig. 1A). These results were replicated in the PD validation subset (n=54), where all 44 features showed significant differences between groups when compared to both the main study control group and the UKB control groups (independent samples t-tests, FDR *q*=0.05). The abnormal features are primarily located in the sensorimotor and visual areas of the cerebral cortex, while the rest are in the cerebellum, caudate nucleus, and putamen (Fig. 1B). These features remained after controlling for FC variability throughout the cortex (Supplementary Fig. 1), demographic factors (Supplementary Fig. 2; Supplementary Table 3), and head motion (Supplementary Fig. 2) (see Supplement for detailed results).

**Figure 1.**
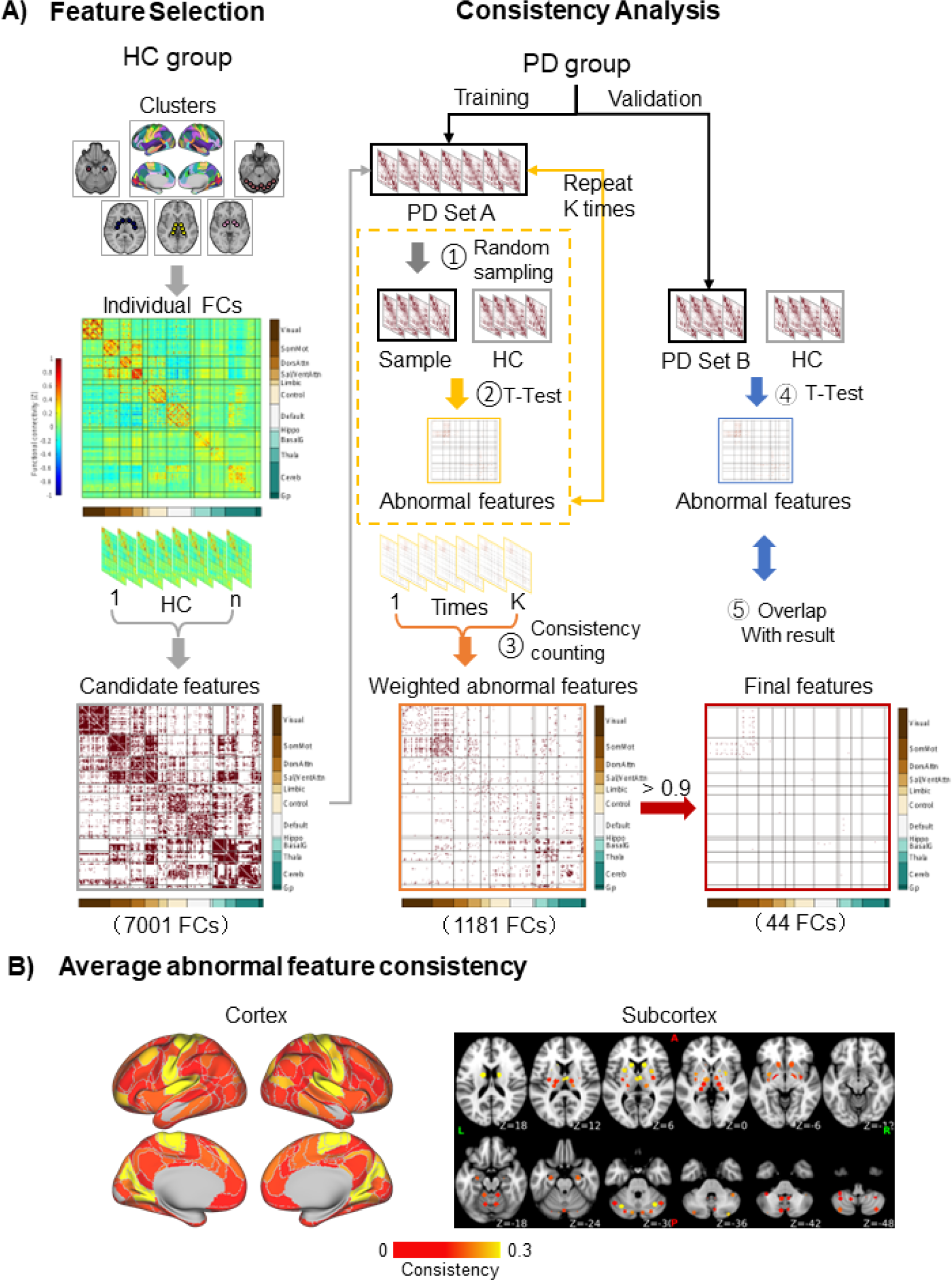
Biomarker identification process. **A)** Feature selection: In the HC group, the FC strengths of all cluster pairs were calculated. Those that were significantly different from 0 (one sample t-tests) were retained, yielding 7,001 candidate features. Consistency analysis: One hundred permutations were performed, in which 51 PD patients were randomly selected from the discovery subset (n=112), forming PD set A (step 1). From the candidate features, abnormal features were found by identifying the features that significantly differed (independent samples t-tests, False Discovery Rate *q*=0.05) between PD patients and HC in each permutation (step 2). This process yielded 1,181 abnormal features. The consistency was calculated for each abnormal feature, defined as the percentage of permutations in which the feature was significantly different between groups. An abnormal FC matrix was generated, weighted by consistency (step 3). The 44 features that were significantly different between groups in ≥90% of the permutations were deemed to be consistently associated with PD (final features). The final features were validated in the independent validation subset of patients, PD set B, by identifying abnormal features (step 4) and determining how many of the 44 final features were among them (step 5). **(B)** The average consistency is shown for each cortical and subcortical cluster. The abnormal features are primarily located in the sensorimotor and visual areas of the cerebral cortex, cerebellum, caudate nucleus, putamen, and thalamus. FC=functional connectivity; HC=Healthy control; PD=Parkinson’s disease.

The FC strength distributions of the 44 final features (Fig. 2A, left) were significantly lower in the PD group compared to controls (K-S test, statistic=0.64, *p*<0.001). This was specific to the 44 final features, as the FC strength distributions of 44 randomly selected functional connections (1,000 permutations) were not significantly different between groups (K-S test, statistic=0.10, *p*=0.69; Fig. 2A, right). The 44 functional connections and their strengths in each group are illustrated in circos plots in Fig. 2B, showing markedly stronger FC strength in the controls.

**Figure 2.**
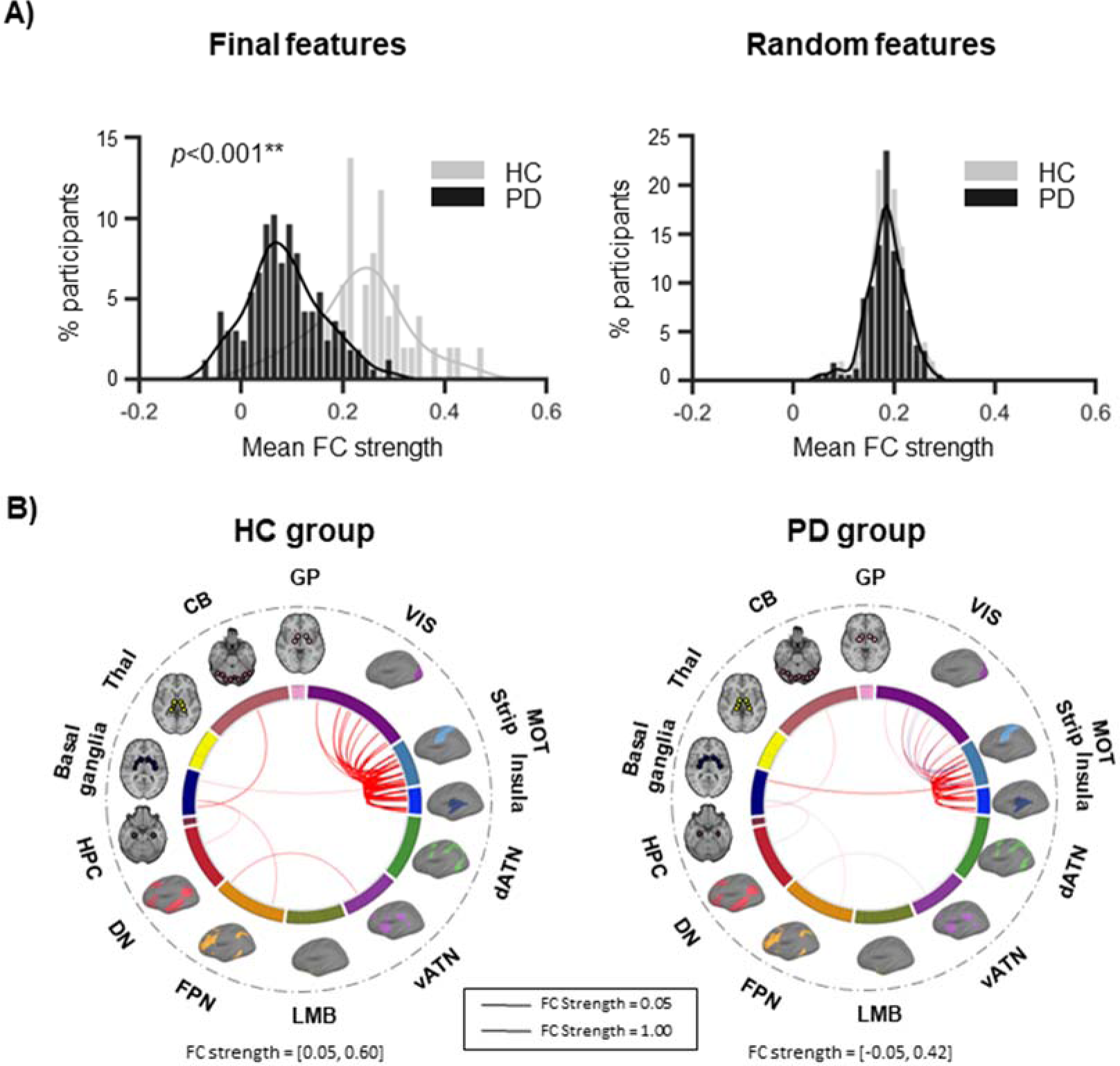
The FC strength of the identified biomarkers is lower in patients compared to HC. **(A)** We plotted the distribution of mean FC strength, binned into intervals of 0.015, for the 44 final features in the whole PD group and in the HC group. The two distributions were significantly different (K-S test, statistic=0.64, *p<*0.001), with patients exhibiting lower FC strengths than controls. This was specific to the 44 final features, as the mean FC strength distributions of 44 randomly selected functional connections did not show a significant difference between groups (K-S test, statistic=0.10, *p*=0.69). Shown here is the distribution of 44 random features in one of the 1,000 permutations performed. **(B)** The 44 functional connections are illustrated in each group using circos plots. Red indicates positive connections, while blue indicates negative connections. The width of the lines represents the FC strength. The sensorimotor strip and insula networks are shown separately here for reference, however they were grouped in the same large-scale sensorimotor network in the analyses. \*\**p*<0.001 CB=cerebellum; dATN=dorsal attention; DN=default; FC=functional connectivity; FPN=frontoparietal; GP=globus pallidus; HC=healthy control; HPC=hippocampus; LMB=limbic; MOT=sensorimotor; PD=Parkinson’s disease; Thal=thalamus; vATN=ventral attention; VIS=visual.

### Connectivity markers of PD are associated with PD symptom severity

Next, to determine whether the markers capture important information about the pathophysiology of PD on an individual-level basis, we performed multiple linear regressions and LOOCV to predict motor and mood measures (see Table 1 for group means) in all PD patients based on the FC strengths of the markers. The between-network markers (n=33) are mostly comprised of connections between the sensorimotor and visual networks, and within-network markers (n=11) are comprised solely of sensorimotor connections (Fig. 3A). In terms of motor measures, significant correlations were found between the predicted and observed scores for the UPDRS-III (*R*=0.21, FDR-adjusted *p*=0.006), Hoehn-Yahr (*R*=0.22, FDR-adjusted *p*=0.006), tremor (*R*=0.21, FDR-adjusted *p*=0.014), bradykinesia (*R*=0.18, FDR-adjusted *p*=0.032), and upper extremities (*R*=0.24, FDR-adjusted *p*=0.007). There were trends towards significance for rigidity (*R*=0.16, FDR-adjusted *p*=0.058) and gait (*R*=0.15, FDR-adjusted *p*=0.060), but no significant correlation for lower extremities (*R*=0.12, FDR-adjusted *p*=0.120) (Fig. 3B; Supplementary Table 4). In terms of mood measures, significant correlations were found for both the HAMD (*R*=0.23, FDR-adjusted *p*=0.006) and HAMA (*R*=0.21, FDR-adjusted *p*=0.006) (Fig. 3C; Supplementary Table 4). The model coefficients of the between- and within-network markers are illustrated as bar graphs in Fig. 3 and listed in Supplementary Table 4. They show that the between-network markers successfully explain the variance in all motor measures (*p*’s<0.05) except the lower extremities (*p*>0.05), and that the within-network markers successfully explain the variance in mood measures (*p*’s<0.05).

**Figure 3.**
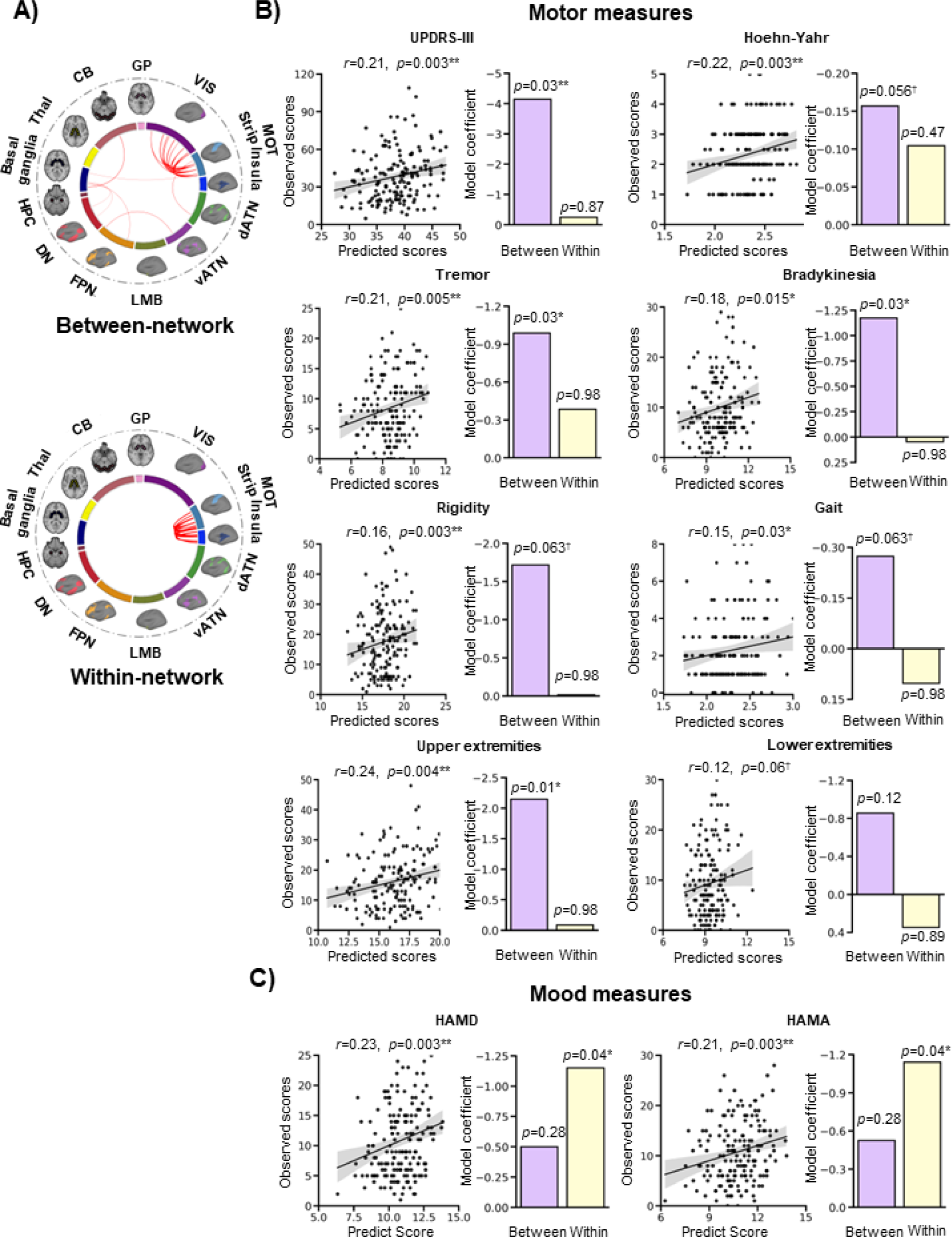
FC markers track with motor and mood symptoms. **(A)** Markers were divided into between-network and within-network functional connections. The sensorimotor strip and insula networks are shown separately for reference, but were grouped together as part of the large-scale sensorimotor network in the analyses. **(B)** Multiple linear regressions were conducted to predict motor and mood measures in patients based on the FC strengths of the within- and between-network markers. Significant correlations were found between the predicted and observed scores for motor measures, including the UPDRS-III (*R*=0.21, FDR-adjusted *p*=0.006), Hoehn-Yahr (*R*=0.22, FDR-adjusted *p*=0.006), tremor (*R*=0.21, FDR-adjusted *p*=0.01), bradykinesia (*R*=0.18, FDR-adjusted *p*=0.03), and upper extremities (*R*=0.24, FDR-adjusted *p*=0.007). The predictions were marginally significant for rigidity (*R*=0.16, FDR-adjusted *p*=0.058) and gait (*R*=0.15, FDR-adjusted *p*=0.060), but not for the lower extremities (*R*=0.12, FDR-adjusted *p*=0.12). The shaded band indicates the 95% confidence interval of the best-fit line. **(C)** The FC markers successfully predicted HAMD (*R*=0.23, FDR-adjusted *p*=0.006) and HAMA (*R*=0.21, adjusted *p*=0.006) scores. The model coefficients of the between-network and within-network markers are shown in the bar graphs. Generally, the between-network markers successfully explain the variance in the motor measures, and the within-network markers successfully explain the variance in mood measures. \**p*<05; \*\**p*<0.01; ^†^0.05>*p*<0.1 CB=cerebellum; dATN=dorsal attention; DN=default; FPN=frontoparietal; GP=globus pallidus; HAMA=Hamilton Anxiety Rating Scale; HAMD=Hamilton Depression Rating Scale; HPC=hippocampus; LMB=limbic; MOT=sensorimotor; PD=Parkinson’s disease; Thal=thalamus; UPDRS-III=Unified Parkinson’s Disease Rating Scale-III; vATN=ventral attention; VIS=visual.

We further explored the contributions of the within-network measures, which consisted in 11 functional connections inside the large-scale sensorimotor network, in the prediction of the mood scales. This network bundles together classic sensorimotor regions as well as a portion of the insula and the auditory cortex. To parse out the contribution of these areas, we divided the network into two: one portion included the classic sensorimotor strip (n=5 connections), and the other included the insula and auditory cortex (n=6 connections) (Fig. 4A). We repeated the prediction analysis for the mood measures using a model with these two subdivisions. There were significant correlations between predicted and observed scores for the HAMD (*R*=0.22, *p*=0.004) and HAMA (*R*=0.21, *p*=0.004) (Fig. 4B). The insula/auditory cortex connections explained the variance in mood measures well (HAMD: *p*=0.08; HAMA: *p*=0.02) while the sensorimotor strip connections did not (HAMD: *p*=0.35; HAMA: *p*=0.86) (Supplementary Table 5).

**Figure 4.**
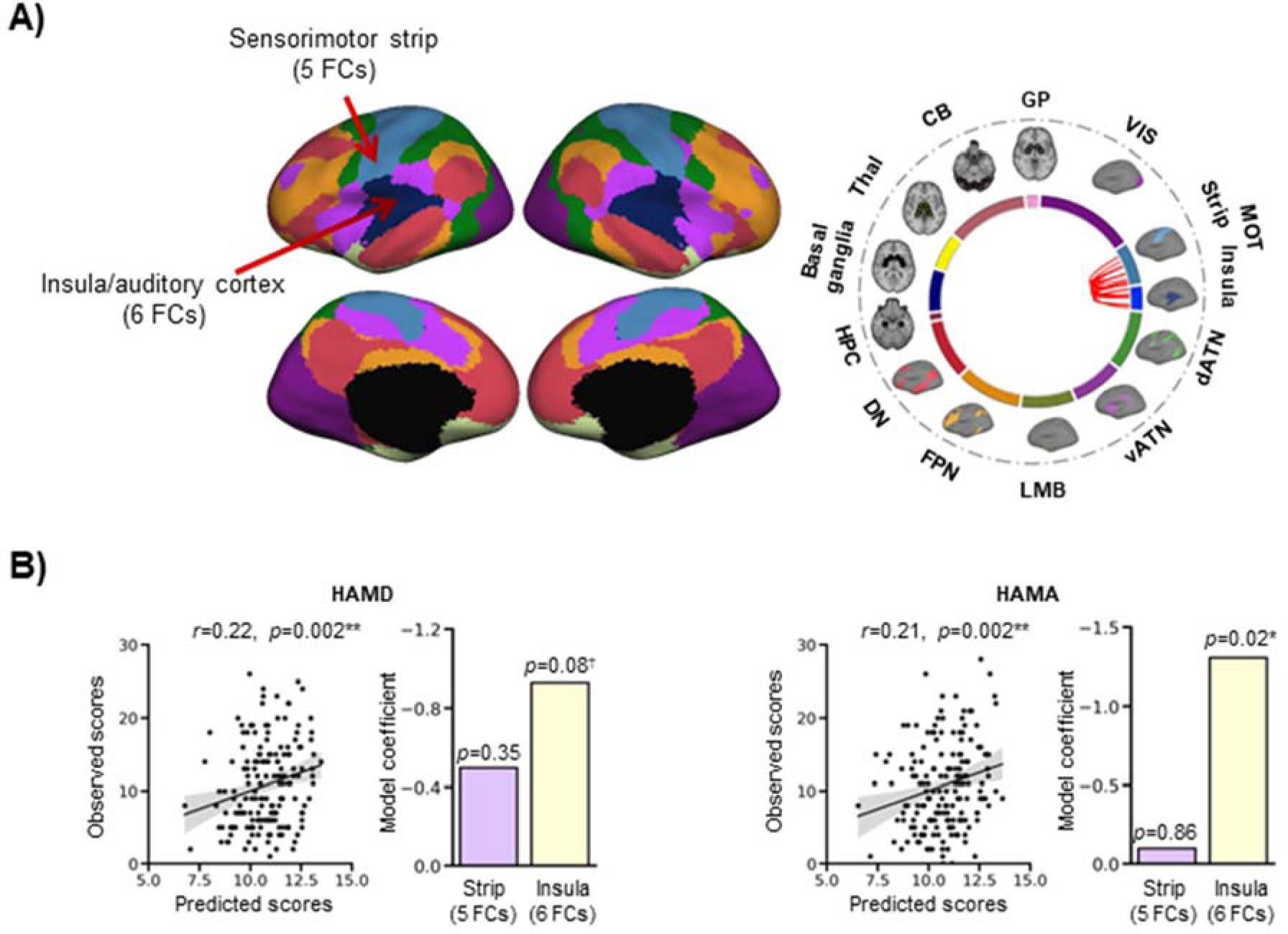
The insula connections are important predictors of mood measures. **(A)** The sensorimotor network was divided into two: one portion included the classic sensorimotor strip (n=5 connections), and the other included the rest of the network, i.e. the insula and auditory cortex (n=6 connections). The circos plot illustrates the within-network connections of the prediction model, all located in the large-scale sensorimotor network. Most of these connections link the sensorimotor strip and the insula/auditory cortex. **(B)** There were significant correlations between predicted and observed scores for the HAMD (*R*=0.22, *p*=0.004) and HAMA (*R*=0.21, *p*=0.004). The bar graphs show the model coefficients of the two portions of the sensorimotor network. The insula/auditory cortex connections contributed to the prediction of mood measures (HAMD: *p*=0.08; HAMA: *p*=0.02), while the sensorimotor strip connections did not (HAMD: *p*=0.35; HAMA: *p*=0.86). The shaded band indicates the 95% confidence interval of the best-fit line. \**p*<05; \*\**p*<0.01; ^†^0.05>*p*<0.1 CB=cerebellum; dATN=dorsal attention; DN=default; FC=functional connectivity; FPN=frontoparietal; GP=globus pallidus; HAMA=Hamilton Anxiety Rating Scale; HAMD=Hamilton Depression Rating Scale; HPC=hippocampus; LMB=limbic; MOT=sensorimotor; Thal=thalamus; vATN=ventral attention; VIS=visual.

### Connectivity markers of PD are generalizable

Ten patients with tremor-dominant PD who underwent MRgFUS thalamotomy in a separate study^26^ were used as an additional independent validation sample (2 women, 8 men; mean age=55.40 years, SD=6.87; Supplementary Table 1). The patients exhibited significantly lower total UPDRS-III scores following the intervention (paired t-test, *t*(9)=4.14, *p*=0.003; Fig. 5A, left), and a trend towards a significant decrease in BDI scores (paired t-test, *t*(9)=2.05, *p*=0.07; Fig. 5A, right). There was a significant increase in the markers’ FC strength from pre- to post-intervention (paired t-test, *t*(9)=3.22, *p*=0.01; Fig. 5B), reflecting the alleviation in symptoms. The PD patients’ FC strengths were different from the main study healthy controls at both timepoints (pre: independent samples t-test, *t*(9)=-5.02, *p*<0.001; post: independent samples t-test, *t*(9)=-2.49, *p*=0.02), showing that some abnormality remained after treatment, and concordant with the fact that patients’ tremor and depression symptoms did not completely resolve. The comparison with the UKB control groups yielded similar findings: the MRgFUS patients exhibited significantly lower FC strength in these markers compared to UKB controls at baseline (pre vs. UKB-55: *t*=-2.96, *p*=0.004; pre vs. UKB-123: *t*=-2.59, *p*=0.01). After the intervention, the patients’ FC strength did not significantly differ from the UKB controls’ (post vs. UKB-55: *t*=-1.14, *p*=0.26; post vs. UKB-123: *t*=-0.81, *p*=0.42) (Supplementary Fig. 3). Our prediction analysis yielded positive correlations between predicted and observed scores, however they were non-significant (UPDRS-III: *R*=0.33, *p*=0.16; BDI: *R*=0.37, *p*=0.11; Fig. 5C). The between-network markers trended towards significantly contributing to UPDRS-III score estimation (*p*=0.054), while within-network markers did not (*p*=0.78) (Fig. 5C, left), consistent with the results of the main study. As expected, the opposite was true for the mood measure: the within-network markers trended towards a significant contribution to BDI estimation (*p*=0.057), while between-network markers did not (*p*=0.56) (Fig. 5C, right). Because the training model used a small amount of data (18 datapoints), we reran the UPDRS-III analysis using the training model derived from the main PD cohort to maximize model accuracy. Here, the prediction analysis yielded a significant correlation between predicted and observed scores (*R*=0.54, *p*=0.007), with between-network markers significantly contributing to UPDRS-III estimation (*p*=0.03) (Supplementary Fig. 4).

**Figure 5.**
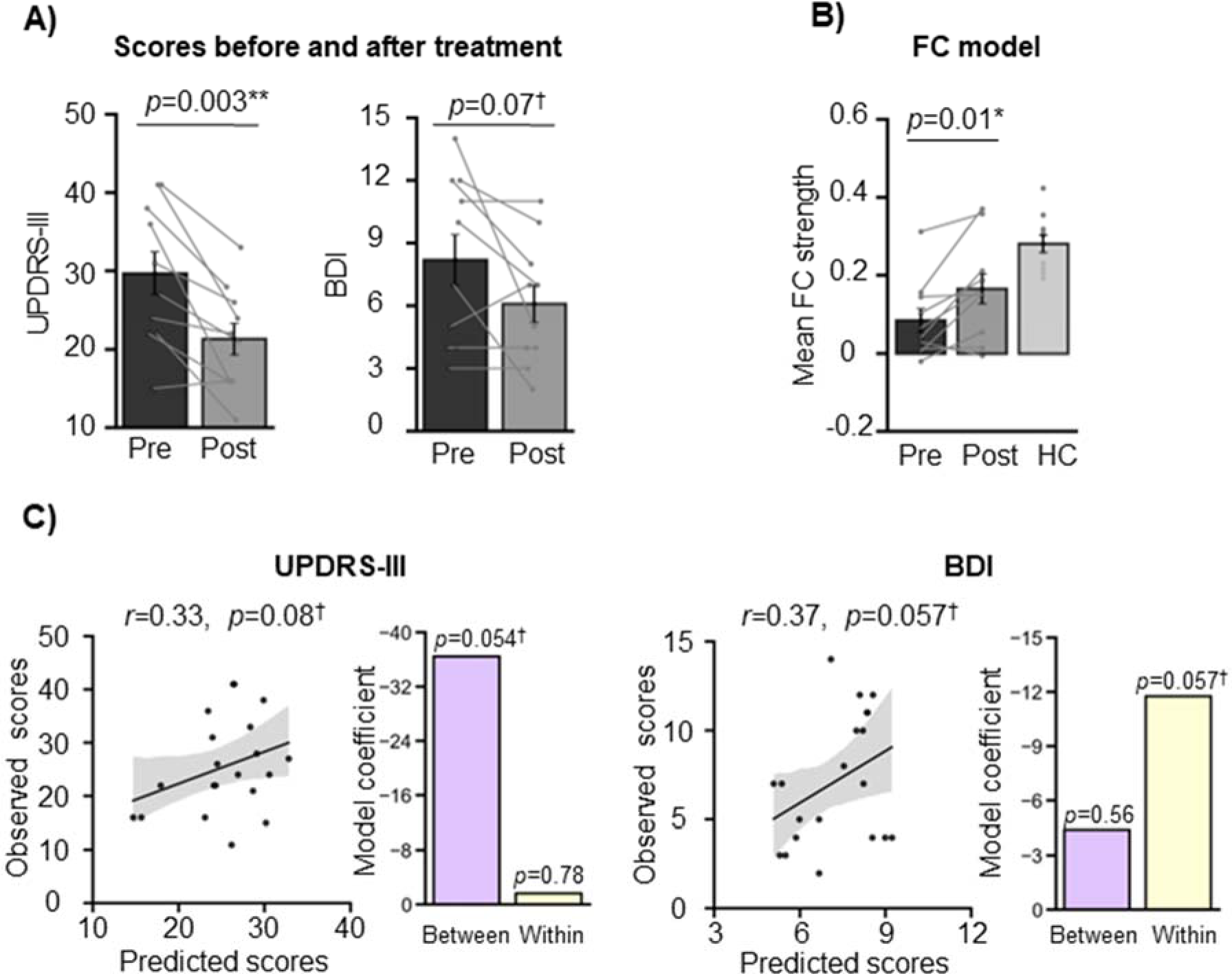
FC marker generalizability in an independent PD dataset. **(A)** In this independent cohort of patients with tremor-dominant PD, MRgFUS treatment successfully alleviated PD symptoms (paired t-test, *t*(9)=4.14, *p*=0.003) and marginally lowered depression symptoms (paired t-test, *t*(9)=2.05, *p*=0.07). Error bars indicate standard errors of the mean and the lines connect datapoints from the same participants. **(B)** Concordantly, there was an increase in the connectivity markers’ FC strength from pre-to post-intervention (paired t-test, *t*(9)=-3.22, *p*=0.01). The PD patients were significantly different from the healthy controls at both timepoints (pre: independent samples t-test, *t*(9)=-5.02, *p*<0.001; post: independent samples t-test, *t*(9)=-2.49, *p*=0.02). The shaded band indicates the 95% confidence interval of the best-fit line. **(C)** Our model was able to predict pre- and post-intervention UPDRS-III and BDI scores, although the predictions were non-significant (UPDRS-III: *R*=0.33, *p*=0.16; BDI: *R*=0.58, *p*=0.007). Between-network connections significantly contributed to the UPDRS-III model (*p*=0.04), while within-network connections did not (*p*=0.91). On the other hand, within-network connections significantly contributed to the BDI model (*p*=0.050), while between-network connections did not (*p*=0.52). \**p*<05; \*\**p*<0.01; ^†^0.05>*p*<0.1 BDI=Beck Depression Inventory; FC=functional connectivity; HC=healthy control; UPDRS-III=Unified Parkinson’s Disease Rating Scale-III.

## DISCUSSION

In the current study, we took an individualized approach to characterize participants’ idiosyncratic functional organization of the brain, through extensive fMRI sampling of patients and individualized brain mapping. This allowed us to identify sets of functional connectivity-based markers that are differentially associated with motor and mood symptoms. Crucially, we demonstrated that these markers are both replicable and generalizable.

### The sensorimotor and visual networks present the most abnormal functional connections

We identified 44 functional connectivity markers that are consistently different in patients with PD relative to healthy control participants, with most belonging to the sensorimotor and visual networks. These connections exhibit reduced functional connectivity strength in patients with PD. The lower FC strength is not a widespread feature of functional brain organization in PD, as randomly chosen functional connections reliably demonstrated equal FC strength across groups. The specificity of the lower FC strength for the identified connections points to a weakening of a network implicated in PD symptomatology. This is in agreement with previous studies that have found disruptions related to PD in the sensorimotor network^3–5, 7, 8, 10–13, 27–30^ and the visual network^4, 8, 10, 12–14, 27, 28, 30, 31^. Although less often studied, the involvement of the visual network is well-documented and is in step with the visual disturbances reported in PD^32^.

### Two sets of functional connectivity markers underlie motor and mood symptoms

The FC markers were found to be associated with total UPDRS-III and Hoehn-Yahr scores, as well as many of the more specific motor symptoms, such as tremor, bradykinesia, and symptoms affecting the upper extremities. The model also did well for the other motor measures, for which the predictions yielded trends towards significance. In addition, the markers successfully predicted depression and anxiety scores. The moderate correlation between predicted and measured scores suggests that the markers provide a neurobiological basis for the motor and mood symptoms of PD, as they can in part explain symptom variability in individuals.

The 44 FC markers we identified consist in 33 between-network connections and 11 within-network connections, and remained robust when controlling for demographics factors and head motion. A majority of the between-network markers connect the sensorimotor and visual networks, while all within-network markers link different regions within the sensorimotor network. These two sets of connections were differentially related to symptoms: the between-network markers contributed most to motor deficit prediction, while the within-network connections contributed most to mood symptom prediction.

Our findings are in agreement with several studies that have investigated the relationship between brain networks and PD symptoms. Motor symptoms are generally associated with dysfunction of the sensorimotor network. For example, Suo et al.^11^ found a correlation between UPDRS-III scores and FC disruption in the postcentral gyrus and the dorsal-most portion of the superior temporal gyrus, which are part of the large-scale sensorimotor network. Koshimori et al.^5^ found that reduced connectivity in the pre-supplementary motor area was correlated with increased UPDRS-III bradykinesia scores. However, some studies found no correlation between motor deficits and network features^10, 12, 13^.

It is interesting that the visual network played such an important role in our prediction model. A minority of our main cohort (22/166 patients) presented with hallucinations and/or psychosis (UPDRS item 1.2), and most of them (15/22) exhibited only slight symptoms. Therefore, it is unlikely that these patients are responsible for the strong contribution of the visual network in our disease pathology model. As mentioned above, however, this finding is not unusual, as several reports found the visual network to be impaired in PD^8, 12–14, 27, 28, 30, 31^, including studies that investigated early PD patients who did not experience hallucinations^4, 10^.

The specific finding that PD is associated with disruptions in sensorimotor-visual network connections has previously been reported. Devignes et al.^8^ reported decreased FC between the sensorimotor and visual networks in PD patients with mild cognitive impairment compared to healthy controls. Lopes et al.^28^ found many functional connections to weaken with worsening cognition in PD (which tends to be associated with older age and more advanced stages of PD), including connections between visual and sensorimotor regions. Wu et al.^30^ identified a pattern of FC associated with greater motor deficits as measured with UPDRS-III scores, which included reduced FC within and between the sensorimotor and visual networks. This between-network disruption could be responsible for the visuomotor deficits observed in PD^33–35^. Taken together, the evidence suggests that it is possible that patients with PD experience visual deficits or vision-related symptoms (e.g., visuomotor) that are not routinely assessed clinically. Additionally, visual hallucinations are a feature of severe, advanced PD, and therefore the presence of visual network disruption in the absence of clinical manifestations of visual deficits may constitute an early sign of future dysfunction.

When it comes to non-motor symptoms, the sensorimotor network, and the middle to posterior insula in particular, played a prominent role in the prediction of depression and anxiety symptoms. The insula as a whole is thought to integrate cognitive, affective, somatosensory, and autonomic signals to guide behavior^36^. The middle and posterior insula seems to especially be involved in somatic and autonomic integration^37, 38^. The HAMD and HAMA scales include many items that concern somatic or autonomic symptoms (e.g., fatigue, gastro-intestinal, cardiovascular, etc.), and the insula FC feature of our model may capture these facets of anxiety and depression. The relationship between mood symptoms and lower FC in the sensorimotor network and insula have previously been reported. Zeng et al.^4^ found significant correlations between mood scores (HAMD, HAMA) and reduced intra-network FC in several networks, including the sensorimotor network, in patients with early-stage PD. Similarly, De Micco et al.^7^ found that PD patients with anxiety had lower intra-network FC in the sensorimotor network compared to those without anxiety. They also observed that lower FC within part of the right insula, which corresponds with the insula/auditory cortex portion of the large-scale sensorimotor cortex, was correlated with greater anxiety severity, as assessed with the Parkinson Anxiety Scale^39^. In Lin et al.^9^, intra- and inter-network FC of the insula was negatively correlated with depression scores (HAMD and BDI). Altogether, the evidence suggests that dysfunction of the sensorimotor cortex, and of the insula in particular, in part underlies depression and anxiety symptoms in patients with PD.

### The FC markers of PD are reliable, replicable, and generalizable

Studies that have sought to discover markers of disease very often suffer from an inability to replicate findings or to generalize their results to cohorts outside the confines of their own experiments. We are aware of only two studies that have investigated FC markers in PD and that attempted to replicate or generalize their findings^15, 30^, and only one was successful^30^. It is therefore paramount that contemporary studies make every effort to replicate their findings as a way to strengthen their results and provide meaningful implications for the field and clinic.

To identify reliable markers, it is crucial to obtain precise fMRI measurements. We and others have previously shown that using individual-level methods to identify functional regions that are unique to each participant allows us to detect effects which are otherwise absent or substantially weaker when group-level methods are used^18–25^. Extensive fMRI sampling of participants is an important step towards capturing individual functional brain organization^40^. In the present study, participants each had 31 minutes of rs-fMRI. In comparison, the aforementioned studies that investigated functional brain networks in PD only acquired 5-10 minutes of data per patient, and used group-level methods.

We demonstrated that our FC markers were reliable by continually comparing our results to chance and minimizing sampling bias by performing numerous permutations of the data and verifying whether the results were consistent and survived thresholding. We tested replicability by identifying the same markers in a separate validation patient subset. Additionally, the fact that our model accurately predicted scores in this subset, and that its prediction ability successfully generalized to an out-of-study cohort of PD patients, substantially strengthens our findings. The use of within-study and out-of-study control groups also served to increase confidence in our results. Finally, our FC markers followed the trajectory of treatment efficacy as their average FC strength increased following treatment. The increase in FC strength cannot be explained by a post-intervention decrease in head motion, as there was no difference in head motion in the pre- and post-intervention fMRI scans (see Supplement). Thus, our findings indicate that our markers may not only explain the pathophysiology underlying symptoms, but also reflect dynamic processes that impact these symptoms.

Taken together, our demonstration of reliability, replicability, and generalizability lends confidence that the FC markers we identified are robust and stable features of PD, and shed light on the pathophysiology of the disease as it relates to motor deficits and mood symptoms.

### Limitations and future directions

First, the PD and MRgFUS patient samples were scanned on the same scanner at the same hospital; it will be important in the future to replicate the findings in an extensively-sampled cohort off-site. Second, the patients in the main and MRgFUS studies were on dopamine replacement therapy. As such, the functional connectivity of these patients may be altered due to long-term medication intake. A future study should seek to replicate the FC markers in a drug-naïve group. Third, the only non-motor scales available in the current study were for the assessment of depression and anxiety. It would be of interest to determine whether the FC markers generalize to other non-motor symptoms.

## METHODS

### Study design and setting

This case-control study was conducted from November 2018 to January 2020, and was performed at Henan Provincial People’s Hospital. Symptom evaluation and neuroimaging were performed at a single time point.

### Participants

#### Parkinson’s disease sample (Dataset I)

We recruited and tested 180 patients with PD. The inclusion criteria included being 18 years of age or older and having a diagnosis of PD according to the United Kingdom Parkinson’s Disease Society Brain Bank criteria^41^. The exclusion criteria were the following: 1) MRI contraindications; 2) history of neurologic disorders other than PD, such as stroke, cerebrovascular disease, seizures, brain tumors, etc.; 3) deep brain stimulation or a prior stereotactic ablation; and 4) average relative head motion ≥ 0.2 mm. Patients were taking one or more of the following medications: madopar, trihexyphenidyl, piribedil, amantadine, entacapone, pramipexole, selegiline, and carbidopa. The average levodopa equivalent dose was 631.08 mg/day. Patients were on medication when they underwent MRI scanning.

#### Control samples

We recruited and tested 62 healthy participants through online advertisement. Screening was performed by phone. Participants had to be 18 years of age or older and devoid of any history of neurologic or psychiatric disorders. They were excluded if they had any MRI contraindications or an average relative head motion ≥ 0.2 mm. Eleven participants were excluded due to excessive head motion.

#### UK Biobank (UKB) control samples

For the replication analyses, we employed two additional control samples from the UKB^42^. 131 healthy control participants of Chinese ethnicity were identified from the UKB database^42^. Inclusion criteria included being at least 18 years old, having Chinese ethnicity, having no history of neurologic or psychiatric disorders, and having a combined 20 minutes of resting-state and task-fMRI data. After excluding participants whose average head motion exceeded 0.2 mm, 123 participants remained (mean age=51.64±6.60; 76 women, 47 men). These participants differed from the PD validation subset in terms of both age (*t*=10.37, *p*<0.001) and sex (*Χ*^2^=5.61, *p*=0.018). To minimize demographic differences, we further selected a subset of 55 participants from the original 123 (n=55; mean age=57.82±3.84; 29 women, 26 men), which significantly differed from the PD validation group in terms of age (although the age was much closer) (*t*=4.84, *p*<0.001) but not sex (*Χ*^2^=1.15, *p*=0.29). When we compared the MRgFUS patients to the UKB control samples, the larger UKB group (n=123) was not significantly different from the MRgFUS group (n=10; 2 women, 8 men; mean age=55.40 years, SD=6.87) in terms of age (*t*=1.71, *p*=0.09), but was significantly different in sex (Fisher’s exact test, *p*=0.02). The UKB subset of 55 participants, however, did not significantly differ from the MRgFUS group in either variable (age: *t*=1.71, *p*=0.09; sex: Fisher’s exact test, *p*=0.09).

#### MRgFUS PD sample (Dataset II)

A referred sample of 10 patients with tremor-dominant PD who underwent MRgFUS treatment at Henan Provincial People’s Hospital were used as an additional independent validation sample (2 women, 8 men; mean age=55.40 years, SD=6.87; Supplementary Table 1). To alleviate hand tremor, the ventral intermediate nucleus of the thalamus (VIM) contralateral to the treated hand was lesioned. The inclusion criteria for the patients were the following: 1) a diagnosis of tremor-dominant PD; 2) postural instability/gait difficulty ratio>1.15 in the medicated [ON] state; 3) tremor at rest≥3 in the affected hand/arm (UPDRS-III item 20), as measured during the medicated [ON] state, or a postural/action tremor≥2 in the hands (UPDRS-III item 21). Patients with bilateral appendicular tremor were included; 4) functional disability due to tremor, as determined by a Fahn-Tolosa-Marin Clinical Rating Scale for Tremor (CRST) score≥2 in any one of items 16-23 in the Disability subsection of the CRST; 5) stable medication for 30 days prior to study entry.

The MRgFUS group and main study control group did not differ in age (independent samples t-test, *t*=-0.11, *p*=0.91) but did in sex (Fisher’s exact test, *p*=0.02). They did not differ in average head motion (independent samples t-test, *t*=-0.31, *p*=0.76). Within the MRgFUS group, there was also no difference in head motion between the two timepoints (paired t-test, t=-0.61, p=0.56).

### Standard protocol approvals, registrations, and patient consents

All participants gave written informed consent according to the Declaration of Helsinki. The study protocol was approved by the Institutional Review Board of Henan Provincial People’s Hospital.

### Outcomes

#### Dataset I

The motor outcomes included the following measures: total MDS-sponsored revision of the Unified Parkinson’s Disease Rating Scale (UPDRS)-III score^43^, Hoehn-Yarn stage^44^, and tremor, bradykinesia, rigidity, gait, upper limb, and lower limb scores derived from the UPDRS-III (see Table 1 for average scores and Supplementary Table 2 for a list of the subitems for each score). Non-motor (mood) outcomes included total Hamilton Depression Rating Scale (HAMD)^45^ and Hamilton Anxiety Rating Scale (HAMA)^46^ scores.

#### Dataset II

The motor outcome was the total UPDRS-III score and the mood outcome was the total Beck Depression Inventory (BDI) score^47^.

### MRI acquisition

#### PD and MRgFUS datasets

Participants underwent one structural MRI scan (8min 50sec) and five resting-state functional MRI scans (6min 14sec; total duration of 31min 10sec). MRI data were acquired on a Siemens 3T Prisma MRI scanner that was equipped with a 64-channel head coil. T1-weighted structural scans were collected using a gradient echo MP2RAGE sequence with the following parameters: resolution=1 mm isotropic, TI1=755 ms, TI2=2500 ms, TE=3.43 ms, TR=5,000 ms, flip1=4°, flip2=5°, BW=240 Hz per pix, echo spacing=7.1 ms, matrix=256×256, 208 slices, with an acceleration factor of 3 (32 reference lines) in the primary phase encoding direction and online GRAPPA image reconstruction. The resting-state fMRI data were acquired using an echo planar imaging sequence with the following parameters: resolution=2.2 mm isotropic, TE=35 ms, TR=2,000 ms, flip=80°, FOV=207 mm, 75 slices. Participants were asked to keep their eyes open, remain awake, and minimize head and body movement. Patient movement was monitored by the MRI operator via a camera. The MRgFUS validation sample underwent the same scans, immediately before and 1 month following the MRgFUS intervention.

#### UKB dataset

MRI data was acquired on a 3T Siemens Skyra scanner equipped with a 32-channel head coil. T1-weighted structural scans were collected using a gradient echo MP2RAGE sequence with the following key parameters: resolution=1 mm isotropic, TR=2,000 ms, matrix=256×256, 208 slices. Two resting-state fMRI scans were acquired using an echo planar imaging sequence with the following parameters: resolution=2.4 mm isotropic, TE=39 ms, TR=735 ms, multiband factor=8, flip=52°, matrix=88×88×64, 490 volumes, with an in-plane acceleration factor of 1. Two task-fMRI scans of 332 volumes were acquired with the same parameters.

### MRI processing

#### Structural MRI

For the PD and MRgFUS datasets, the brain was extracted from the uniform T1-weighted image using the following steps: i) cropping the neck from a gradient echo image (INV2) using FMRIB Software Library (FSL)^48^; ii) extracting the brain using Advanced Normalization Tools (ANTs)^49^; iii) generating a brain mask using ANTs; iv) dilating the brain mask using the Connectome Workbench (https://www.humanconnectome.org/software/connectome-workbench); v) cropping the uniform image using FSL, and vi) applying the brain mask to the uniform image using FSL. Then, in all datasets (including UKB), a surface mesh representation of the cortex was reconstructed using each participant’s structural image, and registered to a common spherical coordinate system^50^. Quality control measures included making sure there were no artefacts, that SNR was adequate, that average motion was under 0.2 mm, and, finally, a functional connectivity sanity check was performed.

#### Functional MRI

The following steps were performed for all datasets: i) removal of the first four volumes; ii) slice-timing correction with Statistical Parametric Mapping 2 (SPM)^51^; iii) motion correction with FSL; iv) registration of fMRI images to the structural image using FreeSurfer^52^; v) normalization of global mean signal intensity across scans using SPM; vi) bandpass filtering (0.01-0.08 Hz) using FreeSurfer; vii) regression of head motion, whole-brain, white matter, and ventricular signals using FreeSurfer; viii) registration to the MNI ICBM 152 T1 asymmetric template; ix) 6-mm full-width half-maximum kernel smoothing using FreeSurfer; and x) projection to fsaverage6 surface space (40,962 vertices in each hemisphere) using FreeSurfer.

### MRI analysis

#### Functional parcellation of the brain

For the PD and MRgFUS datasets, the five fMRI runs were concatenated in each session. For the UKB dataset, the two resting-state and two task-fMRI runs were concatenated. Each hemisphere was divided into five zones (prefrontal, temporal, parietal, occipital, and sensorimotor, the latter consisting of pre-, post- and para-central sulcus regions) using the Desikan-Killiany atlas^53^. We used a k-means clustering approach based on FC profiles to parcellate each zone separately, in order to identify local fine-grained functional networks. To generate connectomes, Pearson correlations were performed between the time series of each vertex and all other vertices. To produce an in individualized parcellation, the cluster boundaries were gradually refined using an iterative approach, described elsewhere^17, 20^. The parcellation generated 152 clusters (76 clusters in each hemisphere). Due to the importance of the cerebellum and several subcortical structures in PD^54^, we additionally applied individual-level parcellations to the cerebellum, thalamus, basal ganglia, and hippocampus using the Seitzman et al.^55^ parcellation, and the globus pallidus using the Pauli et al.^56^ parcellation. The clusters were assigned to one of the seven canonical large-scale functional networks^57^, which included the sensorimotor (MOT), dorsal attention (dATN), ventral attention (vATN), limbic (LMB), frontoparietal (FPN), visual (VIS), and default (DN) networks. This was done by calculating the Dice coefficient between each population-level cluster and the seven networks, and selecting the one that generated the largest Dice coefficient.

### Abnormal functional connections consistently associated with PD

We divided the PD group into two subsets, a discovery subset (n=112; PD set A in Fig. 1A) and a validation subset (n=54; PD set B in Fig. 1A). Using the control group alone, we calculated the FC strength of all the cluster pairs and retained only those whose FC strength was significantly different from 0, using one-sample t-tests. This process yielded 7,001 candidate features, that we then proceeded to compare across the healthy and PD discovery groups using independent samples t-tests with a False Discovery Rate (FDR) *q*=0.05. To mitigate a possible sampling bias, we conducted 100 permutations (random samplings with replacement) to verify whether the significant differences were a product of chance. In each sampling, an equal number of patients with PD from the discovery subset (n=51) and controls (n=51) were selected. The selection was random, and a different set of patients was selected for each permutation. We then measured the consistency of each abnormal feature using a consistency analysis, i.e. the percentage of samplings in which each feature was found to significantly differ between the patient and control groups. The connections that were significant in ≥90% of the samplings were deemed to be stable, abnormal features that are consistently associated with PD (final features). We tested the reliability of the final features in the independent validation subset of patients, testing for significant differences in FC strength against the control group using independent samples t-tests with FDR *q*=0.05. Because the control group was used to compare FC strength against both the discovery and validation subsets, thereby conferring some dependence between analyses, we performed an additional comparison, this time between the validation subset and samples identified from the UK Biobank.

To characterize how the FC strengths differed between groups, we calculated the distribution of average FC strengths of the selected features (binned into intervals of 0.015) in the whole PD group and in the control group, and determined whether the two distributions were significantly different using the Kolmogorov-Smirnov (K-S) test. To ascertain whether the difference in FC strength was specific to the final features or whether it applies to all brain features, we performed the same analysis using 44 randomly selected functional connections, which were permuted 1,000 times.

#### Control analyses

We performed a control analysis to account for variability in FC strength. We calculated the standard deviation for each functional connection in the healthy control group, and regressed out these values from the candidate feature FC strengths. The rest of the procedure was the same as the one described above. To compare these results to the original ones, we performed a Pearson correlation of the consistency of abnormal features between the two sets of results, and calculated the overlap of the selected final features using the Dice coefficient.

Since the patient and control groups significantly differed in age, sex, education, and head motion, we performed additional control analyses to account for these factors. We conducted the same consistency analysis as above using subsets of patients whose demographics were matched, with 100 permutations performed, each of which with a different set of 51 patients. Again, a Pearson correlation of the abnormal features’ consistency was performed between these new and the original results, and the overlap in the final features was calculated using the Dice coefficient.

### Prediction of PD symptoms using abnormal functional connections

A multiple linear regression was performed to model the relationship between the FC strength of the abnormal features and PD symptoms. The FC features were divided into between- and within-network connections, which were entered separately into the model. We used leave-one-out cross-validation (LOOCV) to test the accuracy of the model: the model was trained on the whole PD group minus one patient (n=165), and was used to predict the severity of PD symptoms in the remaining patient (n=1). This process was performed for each of the 166 patients. Pearson correlations were then performed between the predicted and observed scores.

The insula is thought to have an important role in non-motor symptoms of PD^36^. In the large-scale network parcellation, a large part of the insula is included in the sensorimotor network. To determine its role in non-motor symptoms, we conducted a supplementary analysis, where we divided the sensorimotor network into two, one portion comprising the traditional sensorimotor strip, and the other comprising the insula/auditory cortex. We performed a prediction analysis of non-motor symptoms using these subdivisions.

### Validation in an independent dataset

A separate study used MRgFUS to treat hand tremor in 10 patients with tremor-dominant PD^26^. We compared pre- and post-treatment total UPDRS-III and BDI scores, as well as pre- and post-treatment average FC strength across the 44 abnormal features, using paired samples t-tests. We also compared FC strength at each time point to the control group’s using independent samples t-tests. This dataset did not have its own control group, so we used the control group from the main study. To ensure independence, we also performed this analysis using the UKB control groups. Finally, we predicted patients’ pre- and post-treatment scores using the aforementioned method, where 9 patients’ data were used to train the prediction model (18 datapoints), which was then applied to the remaining patient’s UPDRS-III and BDI data to predict their pre- and post-treatment scores. Because this dataset was small, we also used the model trained on the main study PD cohort to predict UPDRS-III scores in the MRgFUS patients.

All statistical tests were performed using scipy v1.6.3. All tests are two-tailed.

### Data availability

The UKB data is available at: https://www.ukbiobank.ac.uk/. The patient and control data that support the findings of this study are available from the corresponding authors upon request.

## Code availability

The code used in this article is available at http://nmr.mgh.harvard.edu/bid/DownLoad.html.

## Acknowledgements

This work was supported by Changping Laboratory (2021B-01-01), China Postdoctoral Science Foundation (2022M720529 and 2023M730175), the Medical Science and Technology Research Project of Henan Province (SBGJ202101002), and NIH grant R61MH121640. L.D. is supported by a Canadian Institutes of Health Research postdoctoral fellowship, FRN: MFE-171291. M.W., H.L., and D.W. have full access to all the data in the study and take responsibility for the integrity of the data and the accuracy of the data analysis.

## Author contributions

Conception and design of the study: L.D., W.Z., J.R., M.W., H.L., D.W.; Acquisition and analysis of data: L.D., Y.B., W.Z., J.R., J.M., W.W., D.W., M.W, H.L.; Drafting a significant portion of the manuscript or figures: L.D., Y.B., W.Z., J.R., S.L., Q.H., X.F., M.W., H.L., D.W.

