## Supplementary for "Individualized functional connectivity markers associated with motor and mood symptoms of Parkinson’s disease"

### Supplementary Results

#### Reliable connectivity markers of PD

Since certain regions, particularly those in association cortex, exhibit greater variability in FC, it is reasonable that areas whose FC is less variable, such as those in the sensorimotor network, would demonstrate more consistency when measuring group differences. To control for this, we repeated the consistency analysis using FC strength variability, measured as the standard deviation in FC strength in the control group, as a regressor. FC strength variability did not affect the consistency of the abnormal features (Pearson $r$=0.99, $p$<0.001), and resulted in the selection of the same 44 final features (Dice=1.0; Fig. S1). We also performed control analyses to account for the significant difference in age, sex, and education between the PD and control groups. We repeated the marker identification process using subsets of PD patients whose demographics matched the controls' on each of these factors (Table S3). Again, the consistency of the abnormal features and selection of the final features remained largely the same (sex: $r$=0.98, $p$<0.001; Dice=0.92; age: r=0.98, p<0.001, Dice=0.92; education: r=0.99, p<0.001, Dice=0.96; Fig. S2). Finally, since the PD group exhibited lower head motion than the control group, we repeated the marker identification process with average head motion as a regressor. The consistencies in the initial and motion-regressed analyses were highly correlated and the final feature selection remained largely the same ($r$=0.99, Dice=0.97).

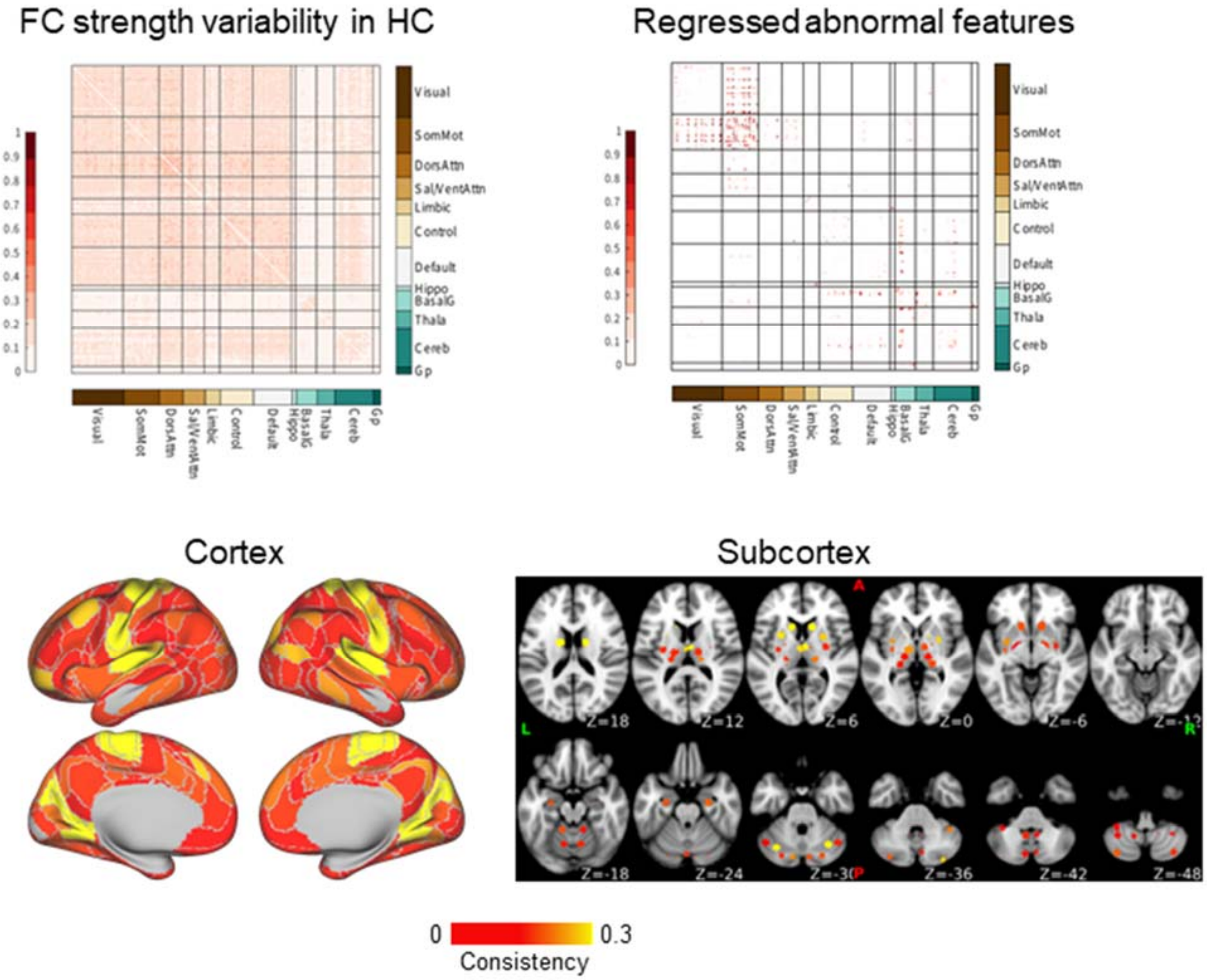


**Supplementary Figure 1 FC strength variability control analysis.** To control for inter-individual differences in FC strength, we repeated the biomarker identification process using FC strength variability as a regressor. FC strength variability did not affect the consistency of the abnormal features, as the clusters' consistency between the initial and control analyses were highly correlated (Pearson *r*=0.99, *p*<0.001). The control analysis yielded the same 44 final features as in the initial analysis (Dice=1.0). The average abnormal feature consistency is shown for each cluster in the cortex and subcortex.
FC=functional connectivity; HC=Healthy control.

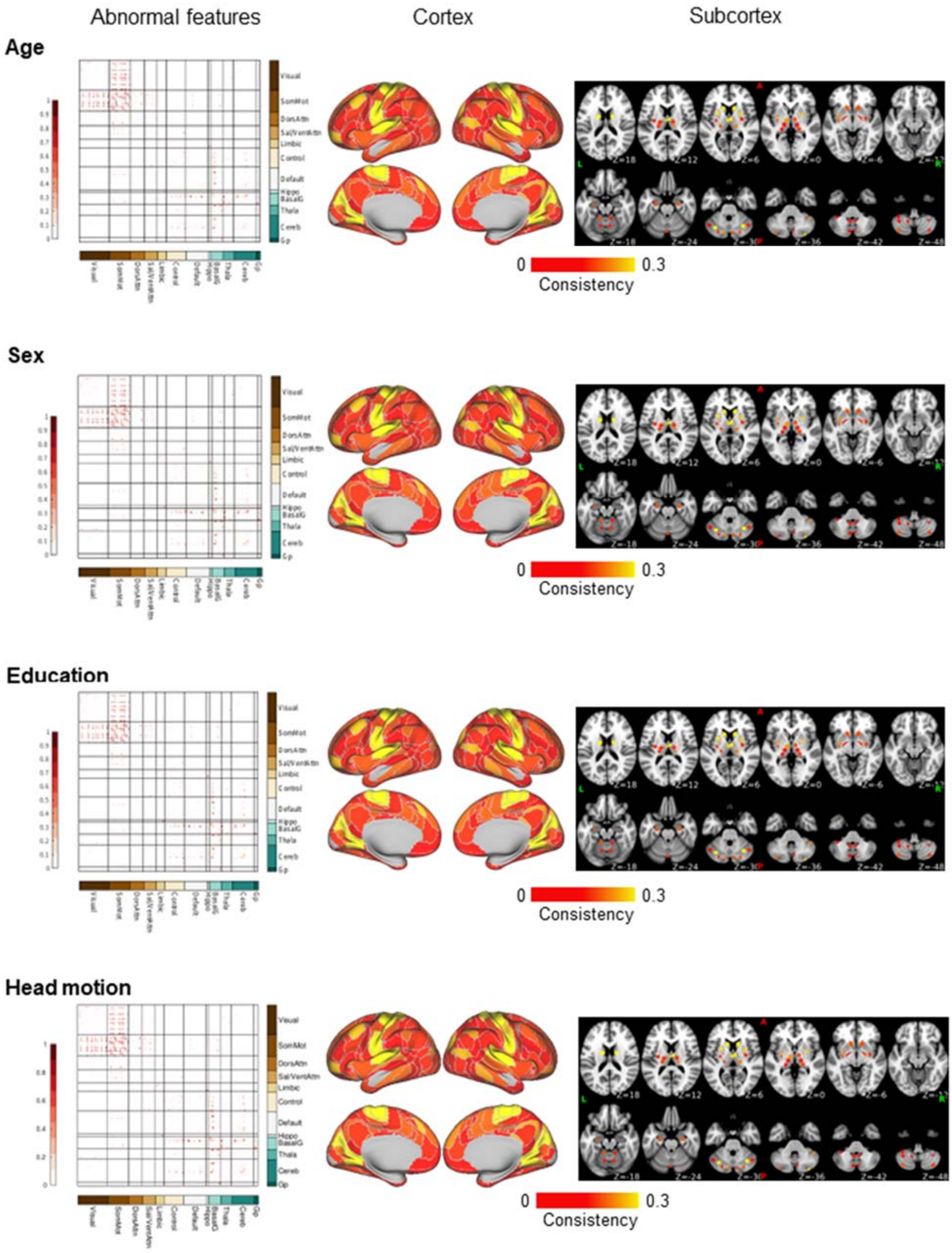
Abnormal features
Cortex
Subcortex
Age
Sex
Education
Head motion
Visual
SomMot
DorsAttn
Sal/VentAttn
Limbic
Control
Default
Hippo
BasalG
Thala
Cereb
Gp
0
0.3
Consistency

**Supplementary Figure 2 Control analysis using demographics and head motion.** We repeated the biomarker identification process within PD subsets that matched the control group in age (1st row), sex (2nd row), education (3rd row), and head motion (4th row). Abnormal feature matrices are shown (left), and the average consistency is shown for each cluster in the cortex (center) and subcortex (right). The consistencies in the initial and demographic- or head motion-matched analyses were highly correlated, and the final feature selection remained largely the same (age: Pearson $r$=0.98, $p$<0.001, Dice=0.92; sex: Pearson $r$=0.98, $p$<0.001, Dice=0.92; education: Pearson $r$=0.99, $p$<0.001, Dice=0.96; head motion: Pearson $r$=0.99, Dice=0.97).

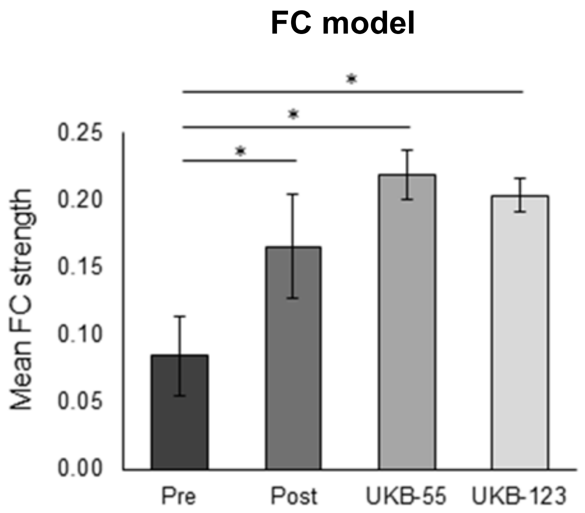


**Supplementary Figure 3 Mean FC strength in the MRgFUS and UKB samples.** At baseline, MRgFUS patients exhibited significantly lower mean FC strength across the 44 FC markers compared to UKB controls (pre vs. UKB-55: t=-2.96, p=0.004; pre vs. UKB-123: t=-2.59, p=0.01). After the MRgFUS intervention, the patients' FC strength did not significantly differ from the UKB controls' (post vs. UKB-55: t=-1.14, p=0.26; post vs. UKB-123: t=-0.81, p=0.42).

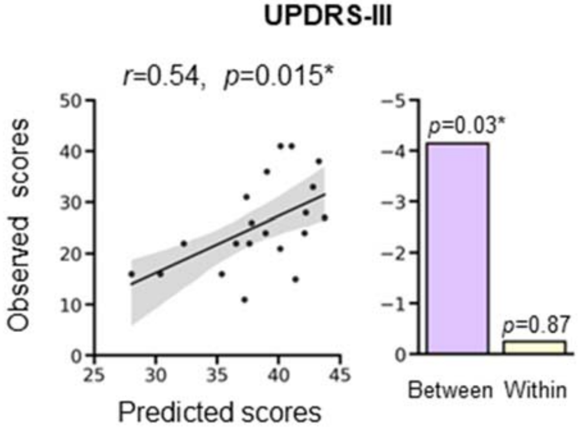


**Supplementary Figure 4 Prediction of UPDRS-III scores in the MRgFUS dataset using the main study model.** There was a significant correlation between predicted and observed UPDRS-III scores (*R*=0.54, *p*=0.007). Between-network markers significantly contributed to UPDRS-III estimation (*p*=0.03), while within-network markers did not (*p*=0.87).

#### SUPPLEMENTAL TABLES

##### Supplementary Table 1. MRgFUS study demographic information

| | MRgFUS |
|---|---|
| Sample size | 10 |
| Sex | 2 W / 8 M |
| Age (SD) | 55.40 (6.87) |
| Years of education (SD) | 10.8 (3.5) |
| Disease duration in years (SD) | 5.20 (1.66) |

MRgFUS=MRI-guided focused ultrasound
SD=standard deviation

##### Supplementary Table 2. UPDRS-III symptom subitems

| Clinical measures | Scale items |
|---|---|
| Tremor | UPDRS-III 3.15, 3.16, 3.17 |
| Bradykinesia | UPDRS-III 3.1, 3.2, 3.6, 3.8, 3.9, 3.14 |
| Rigidity | UPDRS-III 3.3, 3.4, 3.5, 3.7, 3.12, 3.13 |
| Gait | UPDRS-III 3.10, 3.11 |
| Upper extremities | UPDRS-III 3.3 RUE and LUE, 3.4, 3.5, 3.6, 3.15, 3.16, 3.17 RUE and LUE |
| Lower extremities | UPDRS-III 3.3 RLE and LLE, 3.7, 3.8, 3.9, 3.17 RLE and LLE |

UPDRS-III=Unified Parkinson’s Disease Rating Scale-III

**Supplementary Table 3. Demographic information for matched groups**

| K=100 permutations | PD | Control | *p* (SD) |
|---|---|---|---|
| Age-matched | | | |
| Sample size | 51*100 | 51 | - |
| Age (SD) | 58.67 (8.68) | 55.68 (7.62) | 0.07 (0.02) |
| Sex | 16.87 W / 34.13 M | 32 W / 19 M | 0.01 (0.02) |
| Years of education (SD) | 8.76 (3.52) | 10.01 (3.71) | 0.13 (0.11) |
| Sex-matched | | | |
| Sample size | 51*100 | 51 | - |
| Age (SD) | 61.31 (8.10) | 55.68 (7.62) | 0.00 (0.00) |
| Sex | 22.83 W / 28.17 M | 32 W / 19 M | 0.11 (0.06) |
| Years of education (SD) | 8.53 (3.66) | 10.01 (3.71) | 0.08 (0.08) |
| Education-matched | | | |
| Sample size | 51*100 | 51 | - |
| Age (SD) | 61.14 (7.96) | 55.68 (7.62) | 0.00 (0.01) |
| Sex | 18.38 W / 32.62 M | 32 W / 19 M | 0.03 (0.04) |
| Years of education (SD) | 8.87 (3.69) | 10.01 (3.71) | 0.17 (0.14) |

SD=standard deviation

**Supplementary Table 4. Prediction model details**

| Measure | Prediction model *R*, *p*(FDR, 0.05) | Between-network model coefficient, *p*(FDR, 0.05) | Within-network model coefficient, *p*(FDR, 0.05) |
|---|---|---|---|
| Motor | | | |
| UPDRS-III | 0.21, 0.006** | -4.14, 0.034* | -0.24, 0.874 |
| Hoehn-Yahr | 0.22, 0.006** | -0.15, 0.056† | -0.10, 0.190 |
| Tremor | 0.21, 0.014* | -0.98, 0.036* | -0.38, 0.984 |
| Bradykinesia | 0.18, 0.032* | -1.17, 0.036* | 0.04, 0.984 |
| Rigidity | 0.16, 0.058† | -1.71, 0.063† | -0.01, 0.984 |
| Gait | 0.15, 0.060† | -0.27, 0.063† | 0.10, 0.984 |

| | | | |
|---|---|---|---|
| Upper extremities | 0.24, 0.007** | -2.14, 0.012* | -0.08, 0.984 |
| Lower extremities | 0.12, 0.120 | -0.85, 0.127 | 0.34, 0.984 |
| Mood | | | |
| HAMD | 0.23, 0.006** | -0.50, 0.282 | -1.15, 0.041* |
| HAMA | 0.21, 0.006** | -0.52, 0.282 | -1.14, 0.041* |

*$p$<05; **$p$<0.01; †0.05>$p$<0.1
FDR=False Discovery Rate; HAMA=Hamilton Anxiety Rating Scale; HAMD=Hamilton Depression Rating Scale; UPDRS-III=Unified Parkinson's Disease Rating Scale-III.

**Supplementary Table 5. Mood measure prediction using only within-network FC markers**

| Mood measure | Prediction model $R$, $p$ | Somatosensory strip model coefficient, $p$ | Insula/auditory cortex model coefficient, $p$ |
|---|---|---|---|
| HAMD | 0.22, 0.004** | -0.49, 0.353 | -0.93, 0.082 |
| HAMA | 0.21, 0.004** | -0.09, 0.863 | -1.30, 0.023* |

*p<05; **p<0.01
HAMA=Hamilton Anxiety Rating Scale; HAMD=Hamilton Depression Rating Scale; UPDRS-III=Unified Parkinson's Disease Rating Scale-III.
